# Two C++ Libraries for Counting Trees on a Phylogenetic Terrace

**DOI:** 10.1101/211276

**Authors:** R. Biczok, P. Bozsoky, P. Eisenmann, J. Ernst, T. Ribizel, F. Scholz, A. Trefzer, F. Weber, M. Hamann, A. Stamatakis

## Abstract

**Motivation:** The presence of terraces in phylogenetic tree space, that is, a potentially large number of distinct tree topologies that have *exactly* the same analytical likelihood score, was first described by Sanderson *et al,* (2011). However, popular software tools for maximum likelihood and Bayesian phylogenetic inference do not yet routinely report, if inferred phylogenies reside on a terrace, or not. We believe, this is due to the unavailability of an efficient library implementation to (i) determine if a tree resides on a terrace, (ii) calculate how many trees reside on a terrace, and (iii) enumerate all trees on a terrace.

**Results:** In our bioinformatics programming practical we developed two efficient and independent C++ implementations of the SUPERB algorithm by Constantinescu and Sankoff (1995) for counting and enumerating the trees on a terrace. Both implementations yield *exactly* the same results and are more than one order of magnitude faster and require one order of magnitude less memory than a previous 3rd party python implementation.

**Availability:** The source codes are available under GNU GPL at <u>https://github.com/terraphast</u>

## 1 Introduction

It is common practice to infer phylogenies on multi-gene datasets. One way to analyze these is to concatenate the data from several genes or entire genomes into one large super-matrix alignment and infer a phylogeny on it via maximum likelihood (ML) or Bayesian inference (BI) methods. Typically, the sites of such an alignment are grouped into a number of p disjoint partitions (e.g., genes) *P*_1_*,…., P_p_.* Each partition is assumed to evolve according to an independent model of evolution and has a separate set of likelihood model parameters (e.g., substitution rates, branch lengths, etc.).

Super-matrices often exhibit patches of missing data as sequence data for a specific taxon might not be available for *all* partitions *P*_*i*_. Such patches occur because a specific taxon might simply not contain a gene/partition or because the gene/partition has not been sequenced yet. In partitioned datasets, patches of missing data can induce an important effect on the likelihood scores of trees. Under specific partitioning schemes, model settings, and patterns of missing data, topologically distinct trees might have exactly the same analytical likelihood score. We say that, a tree topology resides on a terrace, if it has the same likelihood value as another, distinct tree topology. Recognizing such terraces, determining their size, and enumerating all trees on a terrace therefore constitutes an important step when searching tree space but also for post-processing the results of empirical phylogenetic analyses. Final output trees of tree searches can reside on a terrace and thus, represent only *one* of many possible solutions.

The presence of terraces in likelihood-based inferences was first used implicitly by Stamatakis and Alachiotis (2010) to accelerate ML calculations on patchy super-matrices. One year later, the terrace phenomenon was explicitly named and mathematically characterized by Sanderson *et al.* (2011). Some additional properties of terraces, in particular their impact on bootstrap and other support measures were discussed by Sanderson et *al.* (2015). Chernomor et *al.* (2015, 2016) presented full production-level implementations of terrace-aware topological moves for ML tree searches. Finally, Derrick Zwickl developed a python tool called terraphy for detecting terraces that is available at: <u>https://github.com/zwickl/terraphy</u>.

## 2 Implementation

### Interface

The C and C++ interfaces (see <u>https://github.com/</u> terraphast) take as input: a Newick tree string; a binary matrix *M* of size *n* × *p*, where *n* is the number of taxa and *p* the number of partitions and where every row is annotated by a corresponding taxon name, that denotes if data is available or not for species *i* at partition *j*; a bitmask specifying the computation mode (tree on a terrace; number of trees on terrace; enumeration of all trees on terrace); a destination file pointer to potentially print out all trees on the terrace; a pointer to a big integer library object for storing the number of terraces. For the latter we use the GNU multiple precision arithmetic library (GMP) by default. As we show in our experiments the number of trees on a terrace can easily exceed the 64 bit unsigned integer range. Thus, using GMP is mandatory to prevent integer overflow. The interface function returns an integer that either contains an error code or indicates a successful invocation.

### Limitations

As the library calculates the number of unrooted trees on a terrace given an unrooted, strictly bifurcating input tree, the following limitation applies: The binary input matrix must contain at least one row without any missing data, a so-called comprehensive taxon *tax*_*C*_ such that all p induced per-partition trees *T* | *P*_*i*_ can be consistently rooted on the branch leading to *tax*_*C*_. By induced per-partition tree we refer to the input tree pruned down to the taxa for which sequence data *is* available at a partition *i*. This limitation allows to execute the SUPERB algorithm and, as we show in the supplement, guarantees that the number of rooted trees on the terrace calculated by SUPERB is identical to the number of unrooted trees on the terrace. This limitation can be circumvented by including an appropriate comprehensive outgroup sequence from a reference genome into the dataset.

**Table 1.**
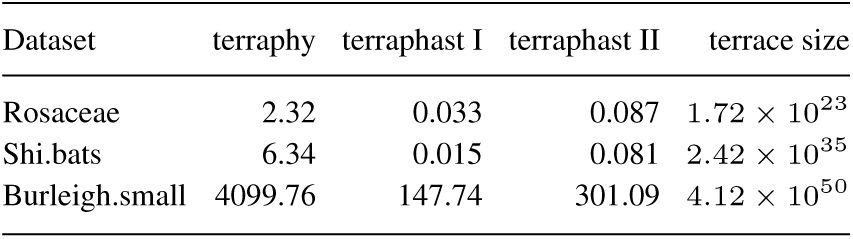
Sequential execution times (seconds) for counting trees on a terrace with terraphy and terraphast I/II.

## 3 Results

We initially tested our implementations on several artificial small 5-taxon datasets for which either all possible trees reside on a single terrace or no terrace exists.

Subsequently, we tested both implementations on 26 empirical datasets from recently published biological studies (available at https://github.com/BDobrin/data.sets) and compared their performance to terraphy. For empirical datasets that did not contain a comprehensive taxon, we sub-sampled partitions such that the samples did contain a comprehensive taxon. For our tests we used a reference system with 4 physical Intel i7-2600 cores running at 3.40GHz and with 16GB main memory. We first verified that our two completely independent implementations (`terraphast I` and `terraphast II`) yield exactly the same results and also compared their run-time performance to `terraphy`. Under identical settings (see supplement for details), all three codes yielded exactly the same number of unrooted trees on *all* datasets, provided that the input tree is rooted at the same comprehensive taxon *tax*_*C*_.

In Table 1 we provide the average sequential execution times over 10 runs and number of trees on the respective terrace for `terraphast I, terraphast II`, and `terraphy` on the three empirical datasets with the largest terraces. All three codes were executed in tree counting mode, that is, enumeration and printout of all topologies on the terrace was disabled. Additional computational experiments under different modes, including memory utilization, parallel performance, and additional empirical datasets are provided in the on-line supplement.

## 4 Conclusions

We have two independent C++ implementations of the SUPERB algorithm for counting trees on a phylogenetic terrace. Because we developed two independent implementations that yield exactly identical results, we are confident that the implementation of the algorithm is correct. Furthermore, `terraphast I` is 28 times faster than `terraphy` on the dataset containing the largest terrace (Burleigh.small) and requires one order of magnitude less RAM (see Table 2 in the supplement). As our experiments with empirical datasets show, a plethora of published phylogenetic trees *do* reside on a terrace. While the phenomenon has been known since 2011, authors of empirical studies do not routinely assess if their tree resides on a terrace. We are optimistic that the availability of an efficient and easy-to-integrate library for this purpose will facilitate integration of this important phylogenetic post-processing step into popular phylogenetic inference tools that are predominantly written in C or C++. `terraphast I` has already been integrated into RAxML-NG (<u>https://github.com/amkozlov/raxml-ng</u>). The authors of GARLI and IQ-Tree also intend to integrate it into their tools.

## Acknowledgements

Part of this work was financially supported by the Klaus Tschira Foundation and DFG grant WA 654/22-2. We thank Olga Chernomor, Bui Quang Minh, and Derrick Zwickl for discussions on the interface definition, Barbara Dobrin for access to her empirical dataset repository, and Alexey Kozlov for integration with RAxML-NG.

## Supplement Section

### Abstract

In this supplement we provide an overview over the SUPERB algorithm. We also show that the number of rooted trees on a terrace as inferred with SUPERB is identical to the number of unrooted trees on a terrace if the unrooted input tree can consistently be rooted on a branch leading to a comprehensive taxon. In addition, we provide details on the test datasets used and discuss some noteworthy implementation details of terraphast I and terraphast II. Finally, we document the C and C++ interfaces and our compressed NEWICK format extension for writing all trees on a terrace to file.

### 1 SUPERB Overview

*Terminology*. First we introduce some terminology and define what the terms we use mean. With ’SUPERB’ we refer to the original algorithm by Constantinescu and Sankoff (1995) for reconstructing supertrees. We use the terms leaf nodes, leaves, and taxa synonymously. We consistently use ’partition’ for subsets of MSA sites that evolve according to the same evolutionary model, whereas ’split’ always refers to a split of leaf nodes (taxa) into subsets. Such splits are denoted by Σ or σ, respectively. The data presence/absence matrix is denoted by M and the comprehensive taxon for rooting by *tax*_*C*_. The induced unrooted per-partition trees are denoted by T|*P_i_,* their rooted counterparts by *T′\P*_*i*_. Finally, the comprehensive input tree is denoted by *T.*

*Original Superb Algorithm*. The original setting of the SUPERB algorithm is as follows: Given a set of rooted binary trees, construct - if possible - all rooted, binary so-called supertrees that are compatible with *all* given trees in the input tree set.

For our purposes, we use all induced per-partition trees as input tree set. These induced per-partition trees *T|P_i_,…, T|P*_*p*_ are extracted from the given input tree *T* (supertree) by pruning all taxa for which no data is available for the specific partition. Therefore, we already know that the algorithm must find *at least* one such supertree. Note that, the input trees of the SUPERB algorithm must be rooted. We describe how the unrooted input trees *T|P_1_,…, T|P*_*p*_ can be consistently rooted in Section 2.

SUPERB consists of two steps: Given the tree set as input, it first constructs a set of constraints that the supertree must fulfill/comply with to fully describe the induced per-partition trees. Then, given these constraints, SUPERB enumerates *all* binary rooted trees that *do* fulfill them.

#### 1.1 Constraint Construction

For the constraint construction, the two children of each node in a tree T| P*i* are ordered (note that, the actual binary input trees are unordered leaf-labeled trees) such that there is a clearly determined left and a right child. The actual order chosen is not relevant as long as it is fixed. By lca(*x*, *y*) we denote the lowest common ancestor of the leaves (taxa) *x* and *y*, that is, the lowest node in the tree that is both in the path from x to the root of the tree and in the path from *y* to the root of the tree. Further, for an inner node x we denote by *x^l^/x^r^* the leftmost/rightmost leaf of x, that is, the leaf we reach if, starting a tree traversal at x, we always descend into the leftmost/rightmost child of a node. The constraints are of the form lca(i,j) < lca(*k,1*). This form denotes that the lowest common ancestor of leafs *i* and *j* must be below the lowest common ancestor (LCA) of leafs *k* and *l* in the supertree we intend to construct. For a given rooted binary tree, it is sufficient to generate one constraint per inner edge (commonly referred to as branches in phylogenetics) (*x, y*), where *x* and *y* are inner nodes of the tree. This constraint for (*x, y*), where *y* is a child of *x* has the form lca(*y^l^,y^r^*) < lca(*x^l^,x^r^*). Note that, depending on whether *y* is the left or right child of *x, y*^*l*^ = *x*^*l*^ or *y*^*r*^ = *x*^*r*^. Thus, due to the symmetric nature of the lca, the extracted constraints are actually of the form lca(*i, j*) < lca(*j, k*). Figure 1 shows a simple example of this constraint construction procedure. In our example, there is only one inner edge (branch) and therefore just one constraint.

**Fig. 1.**
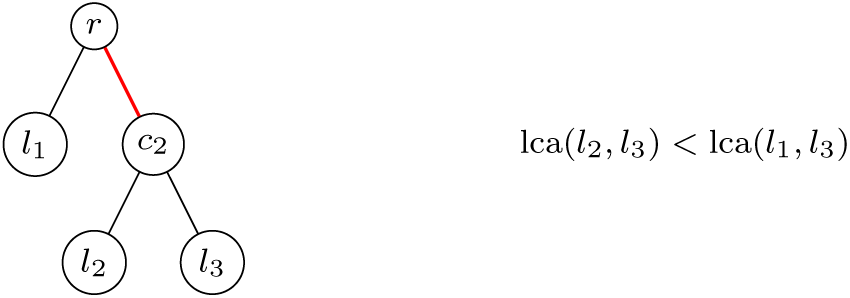
Example tree and the corresponding constraint (for the red edge/branch).

#### 1.2 Tree Enumeration

The main part of the SUPERB algorithm recursively divides the set of taxa/leaves S of the entire tree set, respectively the input tree *T* together with a set of constraints *C*_*s*_ on these leaf nodes. The algorithm starts with all leaves/taxa. Then, for each leaf, it determines if it belongs to the left or right subtree of the root. In the recursion, the algorithm then again divides the leaves among the children of the next node. Therefore, each recursive step corresponds to one node in the supertree we intend to build. The basic insight for dividing the leaves is that for each constraint lca(*i, j*) < lca(*j, k*) the leaves *i* and *j* must be located together in a subtree while *j* and *k* may be separated. Therefore, starting with a trivial splitΣ_0_ = {{*v*}|*v* ∈ Σ} where every node is in its own part, the algorithm iteratively joins for each constraint lca(*i, j*) < lca(*j, k*) the parts *σ*_*α*_ and σ*b* that contain *i* and *j*, respectively. If a supertree exists, at the end of this procedure, there will be a split Σ_*k*_ with at least two parts. If there are exactly two parts, these are the leaves that are located below the two children of the current node. Otherwise, we must consider all possibilities to combine these parts such that we obtain exactly two parts. Each of these possibilities to combine parts results in a different supertree. Thus, enumerating all of them will generate all possible supertrees. For each of the two parts, we call the algorithm recursively with only the leaf nodes in the respective part *and* the constraints that only contain leaves of that specific part. If there are no constraints in one of the recursive calls, it suffices to enumerate all possible binary trees for the corresponding subset of leaf nodes of S. This can be implemented in a straight-forward way.

Let us consider an example with S = {1, 2, 3, 4, 5} and *C*_*s*_ = {lca(1,2) < lca(3,2), lca(4,5) < lca(4,2)}. Then Σ_0_ = {{1}, {2}, {3}, {4}, {5}}. In the next step, we merge 1 and 2: Σ_1_ = {{1, 2}, {3}, {4}, {5}}. Then, we merge 4 and 5: Σ_2_ = {{1, 2}, {3}, {4, 5}}. There are now 3 different ways to split these subsets into two: ({1, 2}, {3, 4, 5}), ({1, 2, 3}, {4, 5}) and ({1, 2, 4, 5}, {3}).

So far, this yields three distinct trees. Now let us consider the recursion into all three partitions:

1. ({1, 2}, {3,4, 5}): For {1, 2}, there are no constraints left and we obtain exactly one tree. For {3, 4, 5} there are also no constraints left. We therefore need to enumerate all possible rooted binary trees with 3 leaves. Those trees all have the form of the tree in Figure 1 and there are exactly 3 of them as we have 3 possibilities for choosing *l*_1_. Thus we obtain 3 trees from this recursion.
2. ({1, 2, 3}, {4, 5}): For {1, 2, 3}, there is the constraint that lca(1, 2) < lca(3,2). Therefore, we join 1 and 2 and obtain ({1, 2}, {3}) asa new split. From the next recursion we obtain exactly one result as there is only one binary tree with two leaves and 1 with one leaf (the leaf itself). For {4, 5} there is also only one tree. Thus, we obtain exactly 1 tree from this recursive step.
3. ({1, 2, 4, 5}, {3}): For {1, 2,4, 5} we have the constraint that lca(4, 5) < lca(4, 2). Thus, we obtain the splits {1}, {2}, {4, 5}. Again, there are three different ways of splitting these subsets into two: ({1, 2}, {4, 5}), ({1}, {2, 4, 5}), ({2}, {1,4, 5}). The first yields exactly one tree from the recursion, the second one also returns only one as the constraint is being used, and the third yields three subtrees as there is no constraint left. Thus we obtain 1 + 1 + 3 = 5 trees from this recursion.

In total, there are 3 + 1 + 5 = 9 possible supertrees for the given set of constraints.

Next, we address the problem of how to root the unrooted input trees for executing SUPERB.

### 2 Rooting by comprehensive taxa

The original SUPERB algorithm is defined on rooted trees. However, all ‘classic’ likelihood- and parsimony-based phylogenetic inference programs and criteria return unrooted trees, except if an outgroup is specified, which however, merely constitutes a drawing option. Therefore, our library function specification explicitly requires a fully bifurcating unrooted phylogenetic tree as input. Depending on the API (Application Programmer Interface) parameter settings, the output is specified to be the number of unrooted phylogenetic trees that reside on the same terrace as the input tree, potentially also including the topologies of all those trees written to a file in NEWICK format.

As the original algorithm is specified on rooted trees we need to devise a method to (i) consistently root our unrooted trees and (ii) ensure that the number of rooted trees reported by the algorithm is identical to the number of unrooted trees. This can be achieved by requiring the input dataset to contain at least one so-called comprehensive taxon *tax*_*C*_, that is, a taxon that *has* data for *all* partitions of the MSA. In other words, the binary input matrix M that contains the presence/absence information about sequence data per species (rows) and partition (columns) needs to contain at least one row that entirely consists of 1s.

If a *tax*_*C*_ exists, every induced tree *T|P*_*i*_ for each partition *i* will contain a branch leading to *tax*_c_. Consequently, each *T*|*P*_*i*_ can be rooted on the branch/edge leading to *tax*_c_ and all induced rooted trees *T′ | P*_*i*_ can therefore be consistently rooted and provided as input to SUPERB.

Given this consistent rooting, we now need to show that the number of trees *and* actual tree topologies rooted at *tax*_c_ as returned by SUPERB are identical to the number of unrooted trees *and* unrooted tree topologies containing *tax*_c_. Without loss of generality, we can order the leaves of the rooted induced trees passed to SUPERB as *tax*_*C*_ first and then all other taxa by some fixed, numerical index (e.g., a lexicographic order).

Given this ordering and rooting the lca(*tax*_*C*_, *y*) will always be the root of the tree for any *T′ | P*_*i*_ and any taxon *y* ∈ *S, y* ≠ *tax*_*C*_ in the tree.

To prove the equivalence of rooted and unrooted supertrees, we show that the mapping `unroot`: 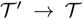 is bijective. `unroot` maps any rooted tree to its unrooted counter-part by simply removing the root. If we restrict the mapping to all supertrees that are enumerated by the SUPERB algorithm, this mapping is surjective onto the set of all unrooted supertrees of the induced subtrees *T|P*_i_. This is because every unrooted supertree is also a rooted supertree of *T′* |*P*_i_ when we place the root into the branch leading to *tax*_*C*_.

However, the mapping is not necessarily injective (see Section 7.1). To ensure that it is injective, we need to restrict the SUPERB algorithm to supertrees where *tax_C_* is a direct descendant of the root node. This can be achieved by fixing the leaf split in the first recursion step such that *tax_C_* is placed in one set and all remaining leaves are placed in the other set. This modification of the SUPERB algorithm does not alter the results and is hence correct based on the following observations: As mentioned above, every constraint involving *tax_C_* is of the form lca(*i,j*) < lca(*j, tax*_*C*_). Thus, *tax*_*C*_ will always be placed into a singleton set when we apply the constraints. Therefore, the fixed split between *tax*_*C*_ and all remaining leaves constitutes a valid split in the original SUPERB algorithm. Since we ignore all other possible splits, the modified SUPERB algorithm enumerates exactly all those rooted supertrees that contain *tax*_*C*_ as direct descendant of the root. Note that, the above modification maintains surjectivity as every tree unrooted at *tax*_*C*_ can be re-rooted at *tax*_*C*_.

Since we have shown injectivity and surjectivity of unroot, the set of unrooted supertrees is equivalent to the set of rooted trees returned by our modified version of SUPERB.

### 3 Implementation Overview

We initially discuss the parts that are common to both implementations before describing some specific implementation details of `terraphast I/II`. Algorithm 1 illustrates the unoptimized pseudo code for both libraries.

#### 3.1 Pre-calculation and Constraint Construction

Before the tree enumerations, our implementations check via function `root_tree` if a comprehensive taxon *tax*_*C*_ exists. If it exists, we root the unrooted input tree *T* on the branch leading to taxon *tax*_*C*_. We unroot and re-root the tree on *tax*_*C*_ if the input tree is given as rooted tree. This defines a fixed traversal order for the rooted tree *T′*. If more than one *tax*_*C*_exists, we select the first valid *tax*_*C*_ that appears when reading M line by line. The SUPERB algorithm is supposed to operate on a set of leaves S extracted by the call to extract_leaves. This step is, depending on the implementation, carried out implicitly by `root_tree`.

After the comprehensive input tree has been re-rooted, we extract all constraints by invoking `compute_constraints`. For each partition P_i_ of the missing data matrix M, we first calculate the induced per-partition tree *T′|P*_i_ via a post-order traversal of *T′* (`extract_partition_tree`). Then, we construct the LCA constraints for each *T′|P*_*i*_ (`extract_constraints`) and combine them into a de-duplicated list.

Duplicate constraints *can* arise when identical subtrees (see Fig. 1 for an example) are induced by more than one partition. Such duplicate constraints can be removed because they provide no additional information on the topology of the *T′|P*_i_. Avoiding unnecessary constraints is another reason for de-duplicating the constraint list. Although SUPERB allows for arbitrary constraints lca(*a, b*) < *lca*(*c, d*), the constraints extracted from the *T′|P*_*i*_ can only be of the form lca(*a, b*) < lca(*b, c*). Note that, SUPERB does not need to distinguish between constraints lca(*a, b*) < lca(b, c) and lca(b, a) < lca(a, c) as they are applied and filtered equivalently. By generating constraints for each *T′P_i_,* we may encounter a specific subtree x times and thus generate *x* instances of the constraint lca(*l*_2_,*l*_3_) < lca(*l*_1_,*l*_3_).

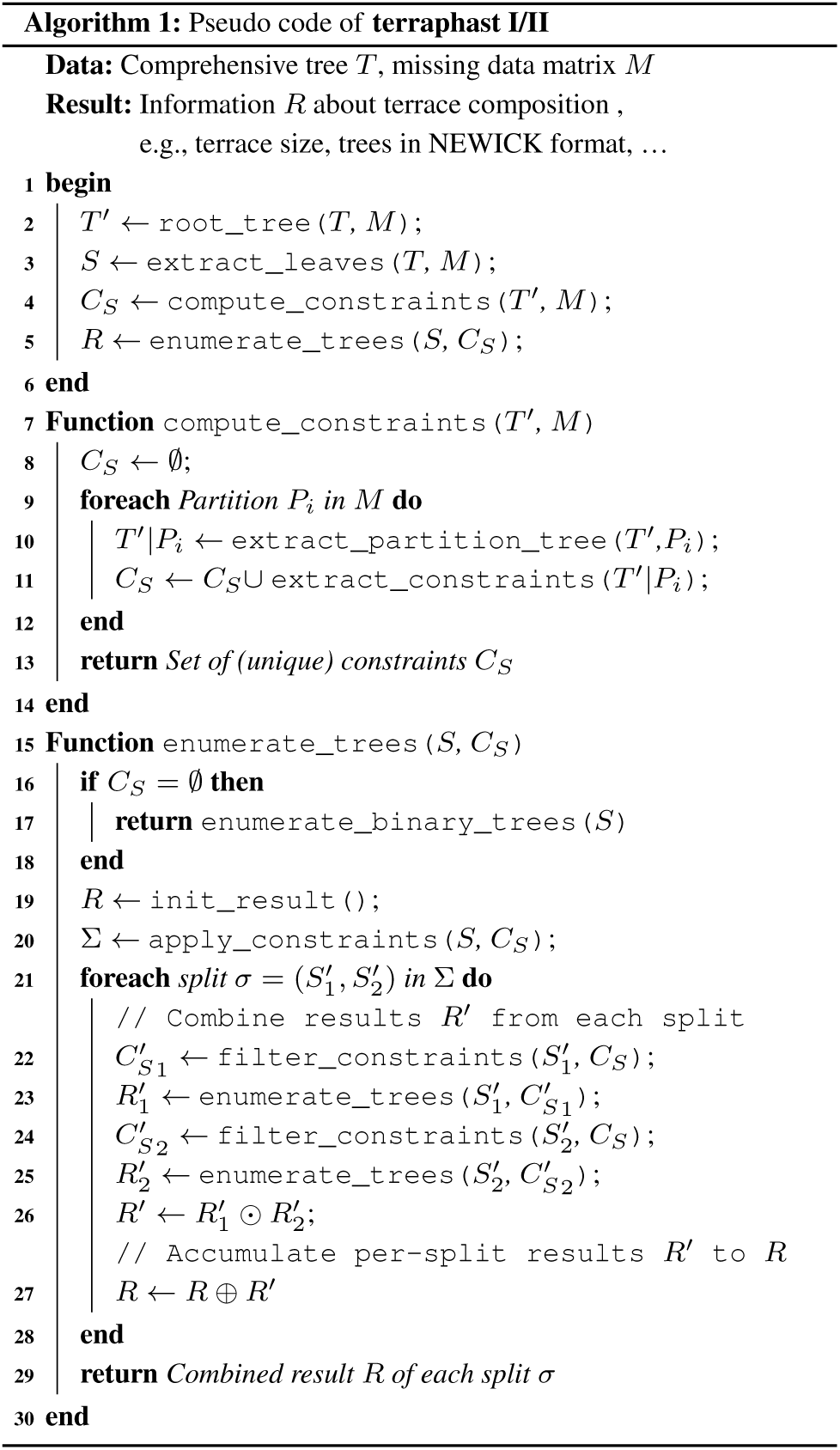

#### 3.2 Tree Enumeration

The function `enumerate_trees` performs the actual terrace analysis. To support various execution modes, this part of our implementations is generic. This means that, function `init_result`, function `enumerate_binary_trees`, operator ⊙s, and operator ⊕s are placeholders for specialized variants of `enumerate_trees`. For instance, if the user wishes to retrieve the exact terrace size, the result to be returned is an integer. In this case, function `init_result` will initialize the result variable *R* as arbitrary precision integer with a starting value of 0, function `enumerate_binary_trees` calculates the number of binary trees for a leaf set *S* if no constraint are left, and the operator ⊙ / ⊕ adds/multiplies intermediate results from the recursive calls. If the user wishes to enumerate all tree topologies on the terrace, all aforementioned functions and operators will generate corresponding tree data structures instead.

Both, `terraphast I` and `terraphast II` provide the following four `enumerate_trees` execution modes:

- *Terrace detection* for checking if the given comprehensive tree *T* resides on a terrace or not.
- Tree *counting* for calculating the number of distinct trees that reside on the terrace.
- *NEWICK tree enumeration* for printing all trees on the terrace to file in NEWICK format.
- *Compressed NEWICK tree enumeration* for printing the compressed NEWICK tree format to file (see Section 8.1)

An important function in the tree enumeration phase is `apply_constraints`, because it accounts for a large fraction of overall runtime. Function `apply_constraints` applies a given set of constraints *S*_*c*_ to a set of sets of leaves generated from the given input set S. It combines, for instance, a set *S*_1_ containing a leaf *l*_*x*_ and a set *S*_2_ containing a leaf *l*_*y*_,*iff*, there is a constraint satisfying lca(*l_x_,l*_*y*_) < lca(*r_x_,r*_*y*_) for any leaf pair *(r_x_, r*_*y*_). Since combining sets, and checking, if a particular leaf is present within a specific set of leaves are frequently invoked operations, they should ideally require constant runtime. To this end, we experimented with two alternative data structures for this task in both implementations. One of them is the union-find with the conventional *union by rank* and *path compression* heuristics. As described by Tarjan (1975), the union-find data structure with both heuristics allows for searching and combining sets in amortized *O*(*α* (*n*)) time where *α*(*n*) is the inverse Ackerman function. This results in an almost constant time operation as *α* (*n*) < 5 for any relevant number *n* < 10^80^ in our context. Alternatively, one can use bit vectors where a set bit indicates if a specific leaf node is contained in the set or not.

After applying the constraints, the algorithm iterates over all possible splits of those leaf sets into pairs of disjunct leaf sets (a specific split) denoted by Σ. For both parts 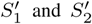 of a particular split, `filter_constraints` determines all constraints 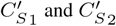 that are still applicable. We then recursively apply `enumerate_trees` on 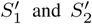 with their corresponding constraint sets 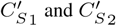 Finally, the ⊙ operation combines the results from both recursive calls, while the ⊙ operation accumulates the results for all possible splits in a single recursive call.

### 4 Implementation of terraphast I

#### 4.1 Constraint Construction

We extract the per-partition trees in a two-step process: For every partition *P*_*i*_, we determine which subtrees of *T′* occur in the induced tree *T′|P*_*i*_. This is calculated simultaneously for all partitions *i* and for every inner node *v* that represents/roots a subtree. More specifically, we compute the bitwise or of the entries in matrix M for both children of v via a post-order traversal. The result is then stored in an augmented presence/absence matrix *M*′.

After extracting the induced per-partition trees, we calculate the constraints as follows. First, we traverse the *T′|P*_*i*_ in post-order to determine the outermost descendant nodes for every node *v*. These outermost nodes have the LCA *v*. We then traverse all edges/branches of all trees *T′|P*_*i*_ once more and use the above information to establish LCA relationships. Constraint calculation is completed by calculating a de-duplicated list of LCA constraints from the above LCA relationships.

#### 4.2 Tree Enumeration

Before the actual tree enumeration, we first use the extracted LCA constraints to construct a so-called multitree. A multitree is a single tree that complies with the constraints and represents all ambiguous (multi-furcating) nodes in the tree via a dedicated node type. Such ambiguities/multi-furcations occur when no constraints exist for a specific subset of nodes. For instance, we represent a three-taxon subtree {a,b,c} as a single node in the multitree. When we generate all possible supertrees based on the multitree, for every possible partial topology not containing {a,b,c}, the {a,b,c} node yields the three subtrees (a,(b,c)), (b,(a,c)), and (c,(a,b)). In other words, a multitree represents the set of all possible supertrees (trees on the terrace) that can be generated from the constraints via SUPERB. A supertree iterator construct (based on regular C++ Standard Template Library iterators) uses this multitree to iterate over, and generate *all* (if the option is set) such supertrees.

#### 4.1 Data Structures

##### 4.3.1 Bitvectors

Subsets of leaves and constraints are represented by packed bitvectors. We implemented an efficient operation to iterate over all set bits in these bitvectors by using bitmasks and the BSF (bitscan forward) instruction. As mentioned above, we calculate the node set of every *T′|P*_*i*_ via a post-order traversal. Here, the bit vector of an inner node is the bit-wise or of its children. The bit vector of a leaf node *i* contains only one set bit at position *i* that corresponds to the respective taxon ID. Our bitvector implementation also provides efficient set operations like union, complement, and symmetric difference. They correspond to the bit-wise operations or, xor, and not which are used throughout our implementation.

In addition, the leaf bitvector is augmented by a constant-time rank support data structure, that is, the rank of an element in the set *l* can be computed efficiently using but a few CPU cycles. The rank operation index only needs to be updated once per recursive call.

##### 4.3.2 Union-find

The union-find data structure uses the *union by rank* heuristic as well as *path compression* to achieve almost constant-time operations. We also implemented an explicit path compression method for the entire data structure to obtain a thread-safe find operation(simple_find). Therefore, we do not need to duplicate these data structures for the threads in the parallel version of the code.

#### 4.4 Memory Allocation

In an early version of terrphast I, memory-management represented the largest performance bottleneck due to the high number of recursive calls and the frequent allocation and deallocation of the respective data structures. Thus, we developed a dedicated memory manager that leverages the LIFO (Last In First Out) structure of the memory allocations in conjunction with the predictable size of the allocations. It maintains a free-list of memory blocks and uses the worst-case (largest) size for every data structure. Note that, the RAM requirements of our implementation are generally low (see Table 2) such that this slight over-allocation of memory is justified.

#### 4.5 Leaf-based Indexing

Since the constraints only contain leaf nodes, we remap all index values (constraints and comprehensive taxon index) from the original indexing based on their node index in the tree to leaf-based indexing. More specifically, we replace every leaf node index by its rank in the leaf set. This allowed us to reduce the space requirements for the leaf bitvectors by a factor of two, and thus further improved spatial locality and, as a consequence, runtime.

#### 4.6 Generalized Implementation

We use an approach similar to the *Strategy pattern* where the SUPERB implementation tree_enumerator relies on a Callback object which implements several callback methods. These callback methods are used for status information (enter, exit, …), execution control (fast_return, continue_iteration), or to provide the elementary operations (combine for ⊙, accumulate for ⊕) and base cases (base_*, null_result,…). Due to static polymorphism, the compiler is able to remove all potential overhead induced by empty method calls.

Analogously to the description in Section 3.2, our implementation provides four different variants of these callback objects: The count_callback and clamped_count_callback callbacks simply count the supertrees. Note that, the results are clamped in case of an integer overflow in the clamped_ variant. The multitree_callback callback constructs a multitree structure that represents all trees on the terrace in a compressed format. Finally, the check_callback callback only checks if there are at least two trees on the terrace. This is accomplished by stopping the split iteration either when the accumulated number of trees is at least two or when a recursive call returns at least two trees (see below).

#### 4.7 Fast Check Heuristic

When checking whether a tree resides on a terrace, we try to prune the recursions of the regular tree counting to the largest extent possible. This is achieved by stopping the iteration over all leaf splits as soon as our accumulated count is ≥ 2. In fact, it is possible to stop the recursion even earlier by using a straight-forward lower bound on the number of equivalent trees generated by a subset of the leaves. Every recursive call returns at least one tree. Thus, the number of possible leaf splits is a lower bound on the number of possible trees at every recursion level. Using this lower bound, we can stop the recursion as soon as we encounter more than two possible leaf splits after the constraints have been applied in a recursive call by invoking the fast_return callback.

#### 4.8 Parallelization

The should_resume_parallel method decides whether the current recursive call should enumerate its different splits in parallel. If this is the case, it prepares all parameters for the recursive calls and aggregates the results after completion of the parallel invocations.

Our parallelization approach could be further improved as follows:

- Currently, all input parameters for the parallel recursive calls are computed in the main thread. By deferring this work to the worker threads, we could potentially further reduce the overhead of the parallel implementation.
- Since the recursive calls often exhibit load imbalance, distributing the workload from two subsequent recursion levels via more sophisticated work-stealing approaches could potentially yield improved load balance.

### 5 Implementation of terraphast II

#### 5.1 Constraint Construction

The function compute_constraints is implemented as described in the Algorithm 1 except that it also omits *tax*_*C*_ and all constraints containing it. This is a valid optimization, because the first iteration of the tree enumeration phase will always generate a split between *tax*_*C*_ and all other leaves. The implementation is aware of the fact, that *tax*_*C*_ only exists implicitly, that is, it is added back after `enumerate_trees` has been completed.

#### 5.2 Tree Enumeration

We use a conventional for loop to iterate over the splits σ. This is achieved by enumerating all possible splits for a specific leaf set/constraint set combination from 1 to 2^*c*-1^ – 1 where *c* is the number of sets that is left after executing `apply_constraints`. The method `get_nth_split` computes the *nth* split by interpreting the number ≥ as a bit vector of length *c*. Each bit *i* set to 1 means that leaf set *c*_*i*_ is supposed to be merged with the set 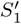. Otherwise, if *i* = 0, the leaf set *c*_*i*_ is merged with set 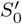 instead. For both parts of the split, `filter_constraints` determines all constraints that are still applicable for the respective part.

#### 5.3 Data Structures

Benchmarks comparing the two alternative implementations suggest that the union-find data structure is on average 24.10% slower in tree counting mode than the bit vector structure. Therefore, `terraphast II` uses bitvectors by default. Users can chose to switch between union-find and bit vectors by editing the header file `leaf_set.h`.

#### 5.4 Generalized Implementation

We implemented the four execution modi of the SUPERB algorithm by using C++ templates and static polymorphism, similar to `terraphast I`. There exists one C++ class for each mode, where each has specialized operator implementations such as `combine_split_results` (⊕ operator) or `combine_part_results` (⊙ operator).

- `CountAllRootedTrees` only counts the number of (sub-) trees by using the arbitrary precision integer type `mpz_class`. The implementation of `combine_split_results`, for example, only calculates the sum over the terrace sizes for individual splits. The recursion stops when no additional constraints can be fulfilled by a leaf set.
- `CheckIfTerrace` determines if *T′* resides on a terrace. This is the case when invoking `combine_split_results` in any step of the recursion yields more than one split. This version of the algorithm stops as soon as a recursive step identifies a terrace for a leaf subset.
- `FindAllRootedTrees` generates all trees residing on the terrace by representing each tree thereon as a corresponding binary data structure. Here, each recursion returns a dynamic array (`std::vector`) containing these binary trees.
- `FindCompressedTree` behaves analogously to version `FindAll RootedTrees`, but only maintains one dedicated data structure that represents the compressed NEWICK tree representation (see Section 8.1) of the terrace.

For each possible split of leaf nodes, the algorithm is then recursively applied to the respective subset of leaf nodes. The intermediate results (terrace detection flag, number of subtrees, or subtree structures) are then combined by `combine_split_results`. The run time contributions and return values of `combine_split_results` vary with the specified execution mode.

#### 5.5 Parallelization

Our parallelization approach is straight-forward and fairly similar to the one used in `terraphast I` (see Section 4.8). The loop, that iterates over all splits returned by `get_nth_split`, is parallelized via the `parallel for` OpenMP pragma. This can be done, because every split represents a different set of trees. Therefore, these iterations are independent of each other. One performance problem is that, the trees from the splits assigned to a thread may be easy/fast to compute, so that this thread finishes long before the others. Hence, parallel efficiency is reduced by load imbalance. To avoid potential overhead by invoking too many threads, the OpenMP parallelization is only applied to the first recursion level at the root. Subsequently, the aforementioned `for`-loop is executed sequentially. As for `terraphast I`, parallel performance could potentially be improved by deploying work-stealing concepts, but is outside the scope of this work.

**Table 1.**
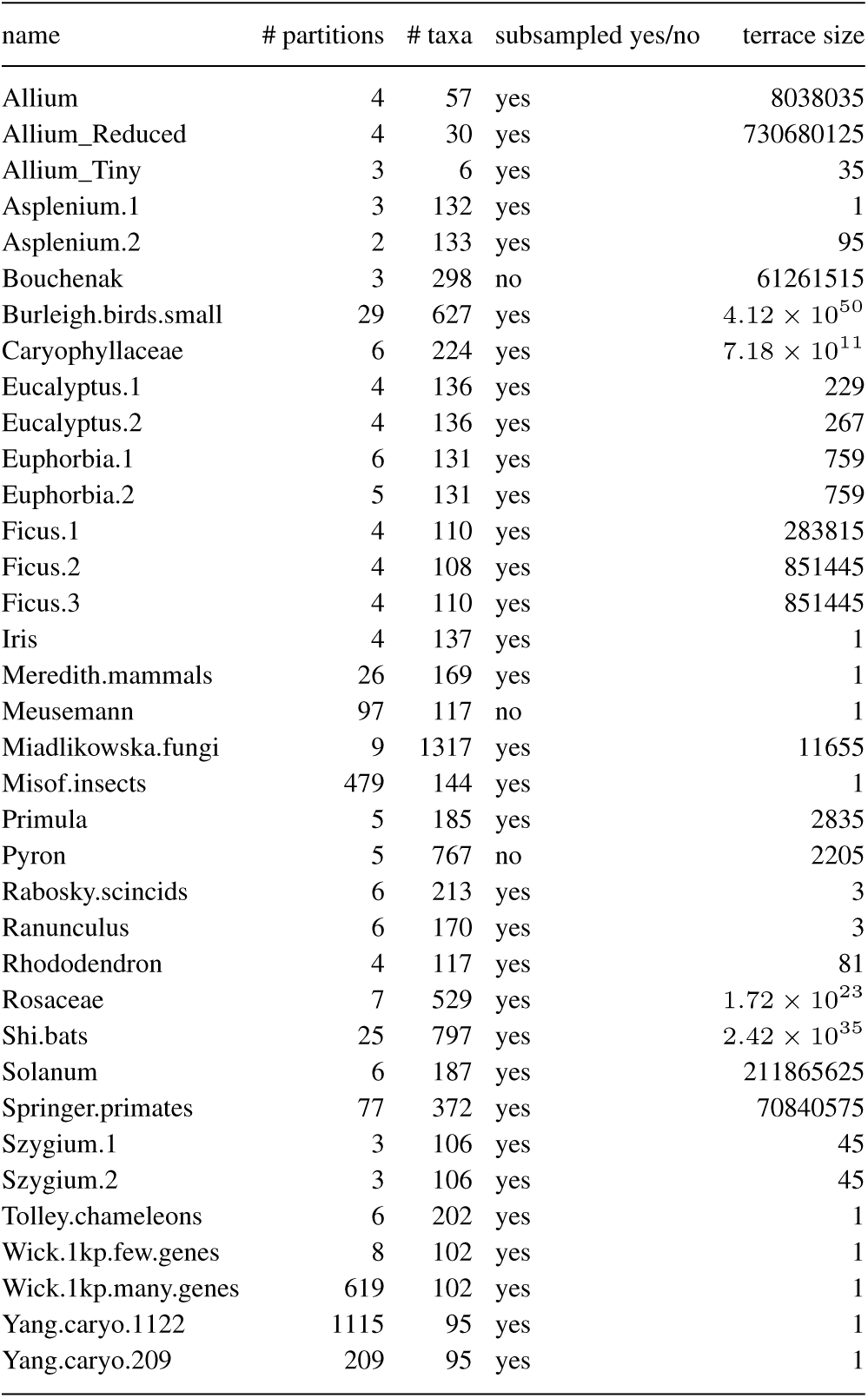
Empirical test datasets used.

### 6 Test Datasets

For testing, we used all empirical test datasets provided at https://github.com/BDobrin/data.sets. The repository contains several recently published partitioned phylogenomic datasets with missing data. As already mentioned in the main text, some of these datasets did not contain a comprehensive taxon *tax*_C_. To this end, we sub-sampled the datasets by applying the following procedure: First, we determined the number of partitions every taxon contains data for. By selecting the taxa with data for the largest number of partitions, we determined candidate taxa which are comprehensive for a large subset of the partitions. We then generated the subsampled data sets by using the cut utility for pruning partitions.

Overall, we generated 36 test datasets from the 26 empirical datasets which are described in Table 1.

In addition, we used some simple artificial datasets for initial testing and verification.

For instance, the following M matrix exhibits a structure where the terrace comprises all possible trees with 5 taxa:

~~~
taxonl 1 0 //taxon1 has data for partition 0 only
taxon2 1 0
taxon3 1 1
taxon4 0 1
taxon5 0 1
~~~

The following M matrix does not exhibit any terraces as there is no missing data:

~~~
taxon1 1 1
taxon2 1 1
taxon3 1 1
taxon4 1 1
taxon5 1 1
~~~

### 7 Additional Experimental Results

In Table 2 we show the memory consumption of `terraphy` and `terraphast I/II` for all test datasets in tree counting mode. Note that, the RAM consumption of `terraphast I/II` is one to two orders of magnitude lower than that of `terraphy`. The larger variance of the RAM consumption in `terraphast I` is due to a memory-wise not fully optimized data structure for storing the induced per-partition subtrees *T|P_i_.* Therefore, this slight waste of RAM becomes more apparent on datasets with a larger number of partitions.

In Table 3 we show run-times for all datasets in terrace detection, tree counting, and tree enumeration modes for `terraphy` and `terraphast I/II`. Note that, terraphy does not offer a terrace detection mode. The results for the tree enumeration mode are incomplete due to excessive run-times.

#### 7.1 Differences between terraphy and our implementations

While conducting our experiments, we noticed that for the Allium_Tiny dataset, terraphy enumerated 37 rooted trees, while there are only 35 unrooted trees on the terrace. This difference stems from the rooting, as in these initial tests the terraphy input tree was *not* rooted at a comprehensive taxon *tax*_*C*_. When we re-rooted the terraphy input tree at a comprehensive taxon, terraphy also enumerated 35 rooted trees which correspond to our 35 unrooted trees due to the consistent rooting.

To further elucidate this, consider the following small example:

~~~
((((s3,s5),s2),s1),(s6,s4));
((((s3,s5),s2),(s6,s4)),s1);
(((s3,s5),s2),(s1,(s6,s4)));
~~~

The three trees above become topologically identical if they are consistently re-rooted at taxon *s*1.

**Table 2.** RAM consumption (MB) in tree counting mode.

| Dataset | terraphy | terraphast I | terraphast II |
| --- | --- | --- | --- |
| Allium | 20.7 | 3.83 | 4.36 |
| Allium_Reduced | 20.3 | 3.80 | 4.27 |
| Allium_Tiny | 19.7 | 3.87 | 4.22 |
| Asplenium.1 | 23.3 | 3.96 | 4.42 |
| Asplenium.2 | 22.8 | 3.89 | 4.34 |
| Bouchenak | 28.5 | 4.01 | 4.71 |
| Burleigh.birds.small | 50.3 | 4.63 | 5.77 |
| Caryophyllaceae | 28.2 | 4.00 | 4.56 |
| Eucalyptus.1 | 23.7 | 3.90 | 4.41 |
| Eucalyptus.2 | 23.8 | 3.94 | 4.78 |
| Euphorbia.1 | 24.2 | 3.93 | 4.27 |
| Euphorbia.2 | 24.3 | 3.89 | 4.81 |
| Ficus.1 | 23.4 | 3.89 | 4.36 |
| Ficus.2 | 23.3 | 3.88 | 8.37 |
| Ficus.3 | 23.3 | 3.91 | 4.38 |
| Iris | 23.7 | 3.96 | 4.43 |
| Meredith.mammals | 61.7 | 3.99 | 4.44 |
| Meusemann | 99.0 | 4.18 | 4.50 |
| Miadlikowska.fungi | 76.4 | 4.68 | 7.52 |
| Misof.insects | 680.1 | 8.24 | 4.47 |
| Primula | 26.0 | 3.98 | 4.46 |
| Pyron | 43.7 | 3.98 | 5.03 |
| Rabosky.scincids | 30.2 | 3.96 | 4.53 |
| Ranunculus | 25.2 | 3.95 | 4.40 |
| Rhododendron | 23.4 | 3.94 | 4.29 |
| Rosaceae | 36.4 | 4.14 | 6.11 |
| Shi.bats | 59.3 | 4.75 | 5.54 |
| Solanum | 25.8 | 4.00 | 4.45 |
| Springer.primates | 130.4 | 5.36 | 4.89 |
| Szygium.1 | 22.8 | 3.91 | 4.35 |
| Szygium.2 | 22.8 | 3.89 | 4.41 |
| Tolley.chameleons | 31.6 | 4.02 | 4.54 |
| Wick.1kp.few.genes | 28.0 | 3.94 | 4.24 |
| Wick.1kp.many.genes | 610.4 | 7.97 | 4.54 |
| Yang.caryo.1122 | 1001.9 | 12.36 | 4.70 |
| Yang.caryo.209 | 210.9 | 5.21 | 4.39 |

*Parallel performance* Finally, in Figure 2 we provide the parallel speedup of `terraphast I` for up to 4 physical cores and up to 8 threads (using hyper-threading) on the reference test system.

The highly frequent memory allocations and deallocations in the algorithm constitute a potential parallel performance bottleneck. To this end, we deployed the dedicated lockless parallel memory allocator `jemalloc` (see http://jemalloc.net/) which yielded up to 25% run time improvement for the GMP-based version that executes a higher number of memory allocations to implement arbitrary precision integers.

In addition, we also assessed if thread pinning, that is, specific thread-to-core assignments, have a notable impact on performance. This is because it is known that thread pinning can substantially affect parallel efficiency on distributed shared memory systems (Klug et al., 2011). In Figure 2 we show speedups for the respective optimal pinning, albeit distinct pinnings did not exhibit a substantial performance impact.

As mentioned before, parallel efficiency could be further improved via appropriate load balancing and work stealing concepts. This was however outside the scope of this work.

### 8 C and C++ Interfaces

In the following we briefly present the C and C++ interfaces.

**Table 3.**
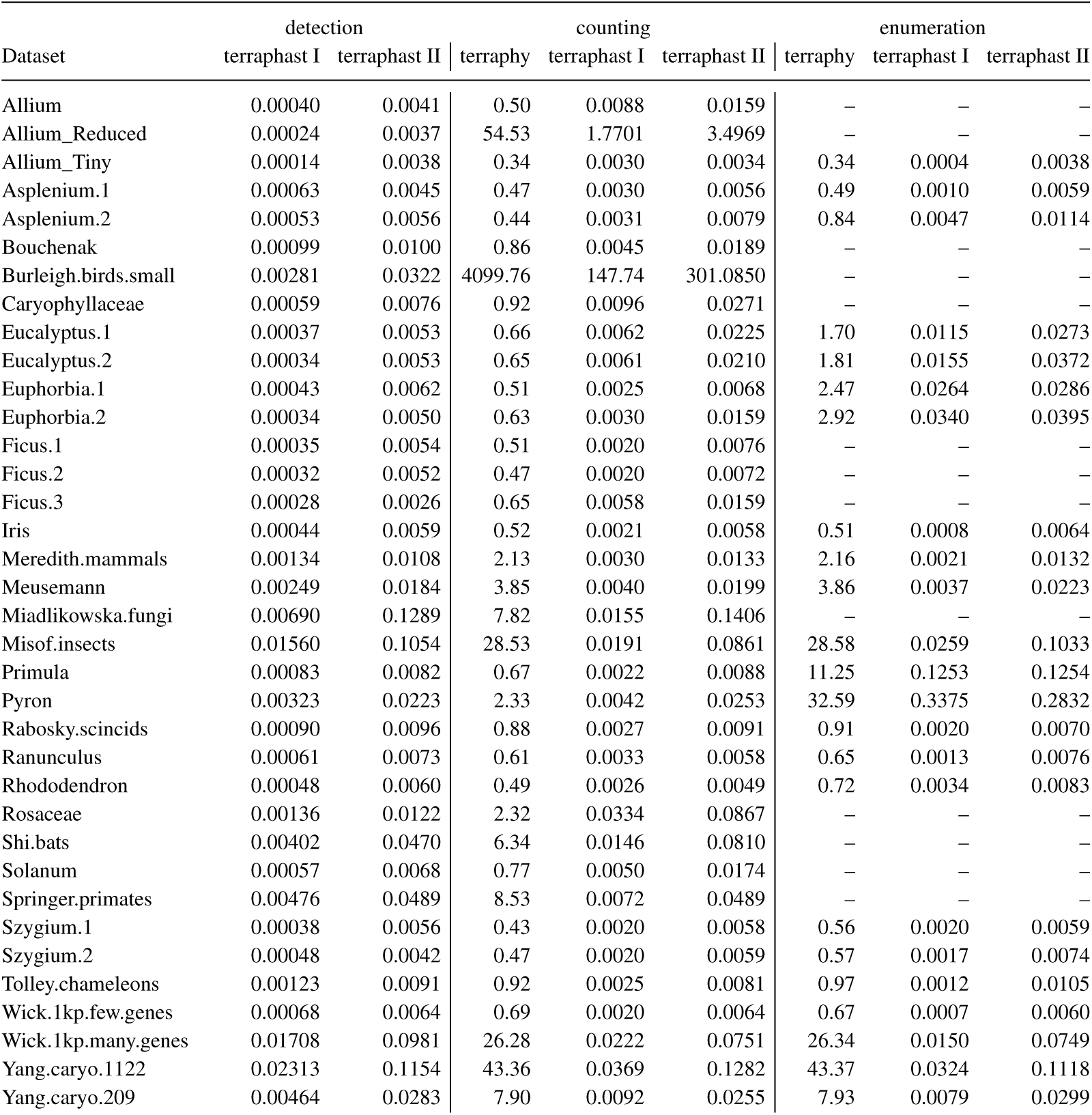
Run times in seconds for terrace detection, counting, and enumeration modes.

**Fig. 2.**
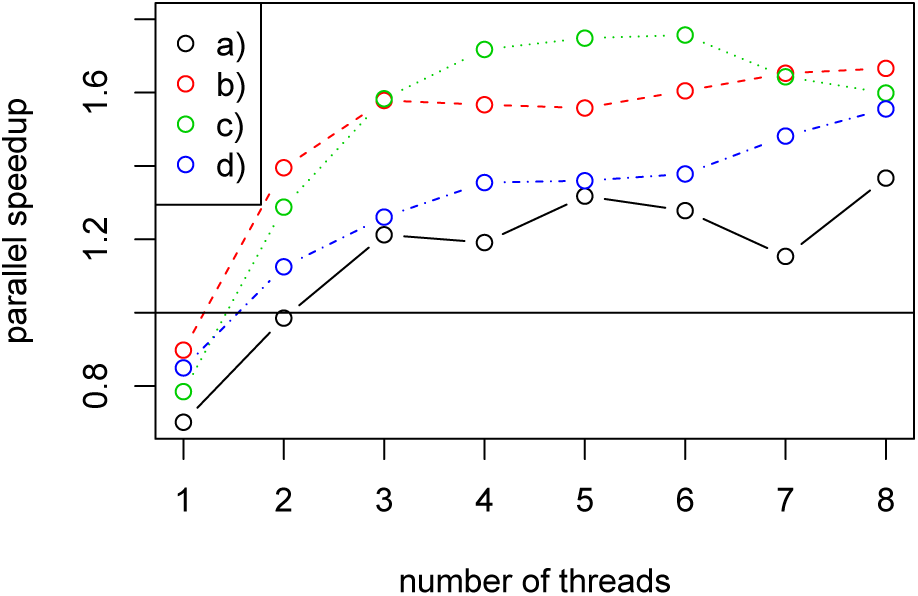
Parallel speedup in tree counting mode for (a) Allium_Reduced with GMP, (b) Burleigh.birds.small with GMP, (c) Allium_Reduced without GMP and (d) Burleigh.birds.small without GMP

*C* interface

~~~
int terraceAnalysis(
missingData *m,
const char *newickTreeString,
const int ta_outspec,
FILE *allTreesOnTerrace,
mpz_t terraceSize);
~~~

Here m represents the binary data input matrix M that also contains a list of taxon/species names for each row. `newickTreeString` is the tree string of the comprehensive tree *T* in NEWICK format that is passed from the application program to the library. The library will then internally compute the induced per-partition trees *T′|P*_*j*_. `ta_outspec` specifies the desired output (execution mode), that is, if the function shall only determine whether the tree is on a terrace or not, if it shall return the number of trees on the potential terrace, or if it shall also enumerate and print to file (in compressed/uncompressed format), all trees on the respective terrace. `allTreesOnTerrace` is a file pointer for printing all trees on the terrace. Finally, `terraceSize` is used to store the number of trees on the terrace, where `mpz_t` is the respective GNU multi-precision library integer type.

**Table 4.**
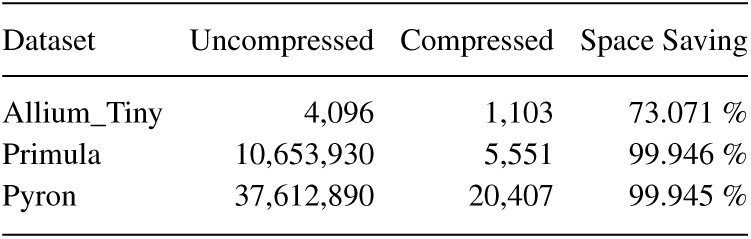
Size of enumerated NEWICK trees in bytes.

| Dataset | Uncompressed | Compressed | Space Saving |
| --- | --- | --- | --- |
| Allium_Tiny | 4,096 | 1,103 | 73.071 % |
| Primula | 10,653,930 | 5,551 | 99.946 % |
| Pyron | 37,612,890 | 20,407 | 99.945 % |

The function returns 0 in case of success and a negative error code to handle errors (e.g., no *tax*_*C*_ could be found, problem parsing NEWICK tree string, mismatch between taxon names in NEWICK tree and m etc.).

*C+ + interface* The C++-interface consists of four families of functions:

~~~
bool is_on_terrace(nwk, matrix)
std::uint64_t get_terrace_size(nwk, matrix)
mpz_class get_terrace_size_bigint(nwk, matrix)
mpz_class print_terrace(nwk, matrix, out)
~~~

The argument-types of `nwk` and `matrix` are `const std::string&` and `std::istream&` (four overloads are provided with all possible combinations). The `out`-argument of the `print_terrace-function` has the type `std::ostream&`. Finally, there is a `*_from_file`-variant of every function-family that takes two filenames as const `std::string&` and reads its data from those files while the other arguments remain unchanged.

If the terrace size exceeds the maximum integer value that can be represented by `std::uint64_t`, the `get_terrace_size` family of functions will simply return the maximum integer value (2^64^ – 1). If the exact terrace size is required nonetheless, the _`bigint-variants` of the functions can be deployed to obtain it. Errors are handled via exceptions.

#### 8.1 Compressed NEWICK representation of a terrace

As enumerating and printing all trees on a terrace to file can easily dominate run-times and require large amounts of disk space, we propose a compressed NEWICK representation that defines all trees on the terrace, but requires substantially less disk space. In Table 4 we provide the compression ratios for output tree files on three representative empirical test datasets. The compressed representation is an extension of the NEWICK format. It relies on the following two extension: First, we use curly brackets to identify a subset of taxa that has no applicable constraint left. For instance, we write {s1,s2,s3} instead of (s1,(s2,s3)); (s2,(s1,s2)); (s3,(s1,s2)); to denote all (rooted) binary trees for these 3 taxa. Second, we use the | symbol to list all subtrees that can be inserted at a specific position in the tree. The expression ((a,(b,c)),(d,(e,f))|(e,(d,f))), for example, is a compressed representation of the two alternative trees ((a,(b,c)),(d,(e,f)); and ((a,(b,c)),(e,(d,f)));. This compressed NEWICK extension could be used, for instance, by tools for post-processing terraces.

